# Sparse Ensemble Machine Learning to improve robustness of long-term decoding in iBMIs

**DOI:** 10.1101/834028

**Authors:** Shoeb Shaikh, Rosa So, Tafadzwa Sibindi, Camilo Libedinsky, Arindam Basu

## Abstract

This paper presents a novel sparse ensemble based machine learning approach to enhance robustness of intracortical Brain Machine Interfaces (iBMIs) in the face of non-stationary distribution of input neural data across time. Each classifier in the ensemble is trained on a randomly sampled (with replacement) set of input channels. These sparse connections ensure that with a high chance, few of the base classifiers should be less affected by the variations in some of the recording channels. We have tested the generality of this technique on different base classifiers - linear discriminant analysis (LDA), support vector machine (SVM), extreme learning machine (ELM) and multilayer perceptron (MLP). Results show decoding accuracy improvements of up to ≈ 21%, 13%, 19%, 10% in non-human primate (NHP) A and 7%, 9%, 7%, 9% in NHP B across test days while using the sparse ensemble approach over a single classifier model for LDA, SVM, ELM and MLP algorithms respectively. The technique also holds ground when the most informative electrode on the test day is dropped. Accordingly, improvements of up to ≈ 24%, 11%, 22%, 9% in NHP A and 14%, 19%, 7%, 28% in NHP B are obtained for LDA, SVM, ELM and MLP respectively.

## I. Introduction

Intra-cortical Brain Machine Interfaces (iBMIs) have made significant strides from the first neuronal population level studies reported in non-human primates (NHPs) [1], [2] to the recent impressive real-time demonstrations in humans [3]–[5]. The aim of iBMIs is to substantially improve the lives of patients afflicted by spinal cord injury or debilitating neurodegenerative disorders such as tetraplegia, amyotrophic lateral sclerosis. These systems take neural activity as an input and drive effectors such as a computer cursor [4], [6], wheelchair [7] and prosthetic [3], [8], paralysed [5] limbs for the purposes of communication, locomotion and artificial hand control respectively.

Despite these compelling advances, barriers to clinical translation remain. One of the most pressing issues is the non-stationary nature of neural recordings (Fig. 3). Recording condition changes such as micro-motion of electrodes (Fig. 1), development of scar tissue, changes in electrodes impedance are prominently responsible for the exhibited nonstationarity across time [9]–[13]. Due to these changes, a decoder trained at a given point of time is often times rendered unusable after tens of minutes to a few hours [11], [14]–[16].

**Fig. 1.**
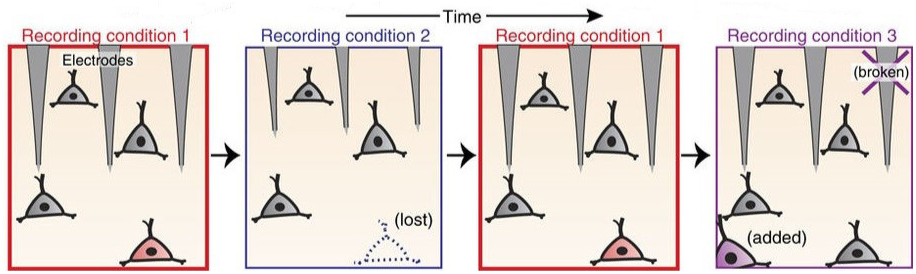
Micro-motion of an electrode array leads to appearance/disappearance of a neuron’s signals at a particular electrode as seen in recording conditions 1,2. Furthermore, in a more dire scenario as seen in recording condition 3, certain electrodes break down and stop yielding any meaningful recorded signal. Figure is not drawn to scale and is adapted from [17] (CC-BY license).

**Fig. 2.**
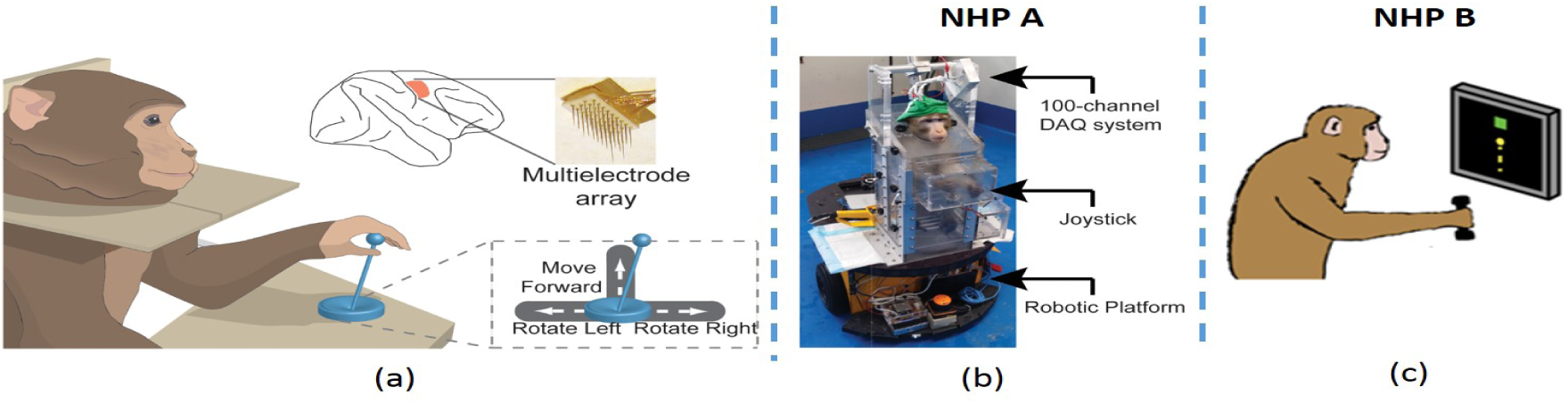
(a) Shows an NHP trained to control a joystick. Joystick movement was restricted to right, left and forward directions only. In the top portion, a multi-electrode array is shown to be implanted in the hand and arm regions of primary motor cortex [19], (b) Experiment 1 setup: NHP A was seated in a robotic wheelchair. The wheelchair translated in the forward, left or right directions in discrete time steps of 100 ms depending on the position of the joystick [19], (c) Experiment 2 setup: NHP B is trained to drive a virtual wheelchair using a joystick in a manner similar to experiment 1. Figures (a), (b) are adapted from [19] (CC-BY license) and Figure (c) is adapted from [20] (CC-BY license)

**Fig. 3.**
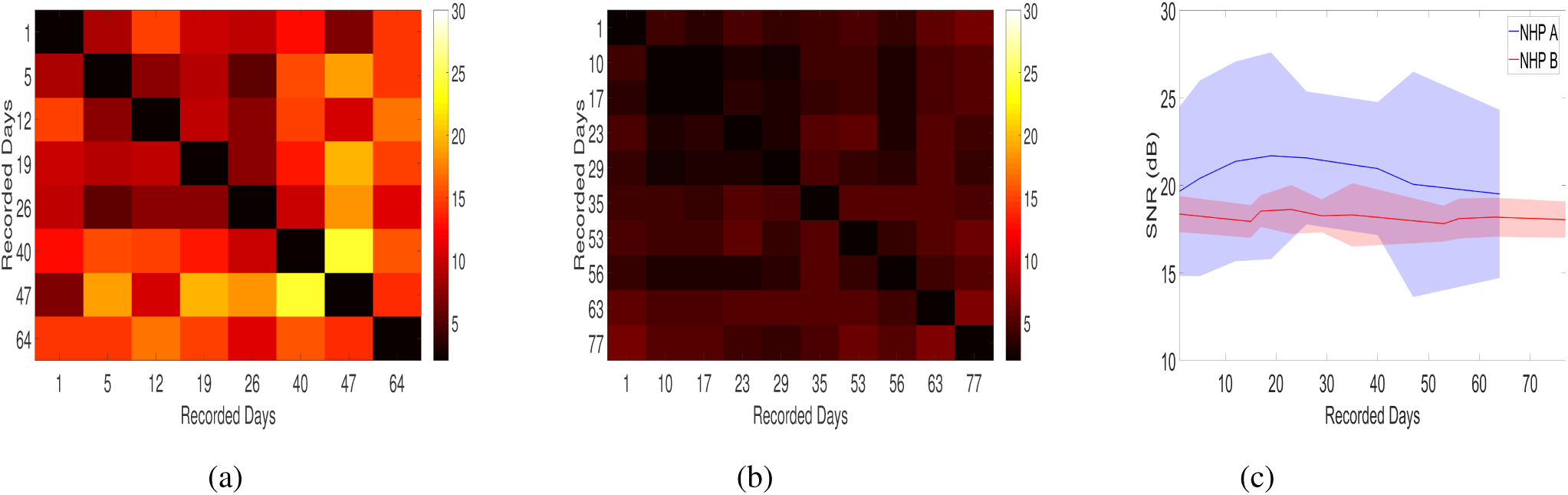
The principal angle is a parsimonious scalar metric to measure the similarity of neural activity across days [17], [30]. (a) and (b) present principal angles between the subspaces of neural activity recorded each day for NHPs A and B respectively. Smaller principal angles denote greater similarity and one can see recorded days being more similar to each other in NHP B than NHP A. (c) presents measured signal to noise ratio (SNR) values following the methodology given in [31] with mean (solid line) and standard deviation (shaded region) values across electrodes plotted for recorded days in NHPs A and B. As far as signal stability is concerned, we can see the average SNR values across days to be stable across days for both NHPs. NHP A shows greater variance in the SNR values among its electrodes compared to NHP B, showing that some electrodes capture better spikes than the others.

To counter nonstationarity, the most commonly employed approach is to re-train the decoder every day using a set of supervised trials at the beginning of a recording session. The subject is made to perform a set of calibration trials and target labels are collected along with neural activity [10], [15], [16], [18]. This approach leads to an increased setup time and necessitates the presence of an expert to get the system up and running. Data collected from previous days/sessions is discarded and the decoder utilizes only the most recently collected data to learn the mapping between neural activity and behaviour.

Despite the deleterious phenomenon of nonstationarity, studies have shown some commonality exhibited between neural recordings across time [11], [21]–[25]. Accordingly, there are alternate approaches that aim to utilize this similarity to build robust iBMIs. For example, researchers in [26] have proposed a principal component analysis based domain adaptation (PDA) technique to reduce decoder calibration times by utilizing historical data adapted with a small sample set of current day’s data. However, consistent decoding results have only been reported for a maximum of three day period of separation between training and testing days and it remains to be seen if this technique still holds over a longer period of separation. Besides, it still resorts to explicit collection of supervised trials data in the beginning of a recording session necessitating suspension of decoder use even if that means for a relatively lesser time than standard calibration times for training a de novo decoder.

In order to avoid suspension of decoder use, an alternate strategy is to update the decoder in an unsupervised manner [14], [24], [27]. These techniques rely on tracking and updating changing neural statistics while the decoder is still in use. Authors in [14], [27] have presented a decoder being updated in a retrospective fashion after every block of trials in a human subject based point and click cursor experiment. Sustained performance has been reported employing this approach in a closed loop study involving 11 sessions spanning 29 days [27] and 6 sessions spanning 42 days [14] in two human subjects. However, reported results have shown some manual intervention with regards to setting of cursor’s speed gain and click thresholds [14]. Furthermore, for a decoder to be updated in a retrospective fashion, there is an implicit assumption about the latest calibrated decoder performing with sufficient accuracy to complete the stated task. [24] reports a self-calibrating method that improves performance over a non-updated decoder in an offline setting in a discrete-control experiment. Results abide by the standard protocol of demonstrating improvements in two NHPs and sustained improvements have been shown to last about 35 and 40 days in two NHPs. However, this approach requires a burn in period [24] before the decoder updates to an optimum level and matches the current day/session’s distribution to yield better results.

Continuing along the lines of eliminating suspension of decoder use, there is another school of thought which focuses on harnessing a large magnitude of past data and builds a data driven robust solution to counter nonstationarity. This approach banks on computationally complex algorithms such as recurrent neural networks (RNNs) [17], long short term memory units (LSTMs), convolutional neural networks (CNNs) [28] to learn complex mappings between heterogeneous neural data across days and effector variables. [17] presents consistent offline performance over 55 and 59 days for two monkeys involved in a hand based target acquisition task employing a fixed multiplicative recurrent neural network (MRNN) as a decoder. Similarly, [28] presents a deep neural network based decoder employing computationally complex LSTM and CNN type neural networks to yield consistent offline results over 381 days in a human subject based study. Although both these offline results are novel and impressive in terms of their sustained performance, the amount of data reported to optimally train these complex models is concerning. Furthermore, these models require hand-tuning of hyper-parameters [17], [28], [29] coupled with specialized hardware during training phase. This makes it perhaps difficult to optimally apply these approaches in a clinical setting.

Taking note of the problems faced by the reviewed approaches, we present a general sparse ensemble decoding approach that performs consistently better than corresponding single classifier models of varying complexity in the face of incoming non-stationary data. Furthermore, we also present data driven hyper-parameter tuning guidelines in *Section* IV-A for different classification techniques to make this approach easier to use.

## II. Materials and Methods

We have based our studies on two different sets of experiments each involving a different NHP. Accordingly, our subsections are split into two parts corresponding to experiments 1 (NHP A) and 2 (NHP B). Two adult male macaques (*Macaca fascicularis*) have been used in the experiments. All conducted animal procedures abide by the standards of the Agri-Food and Veterinary Authority of Singapore and the Singapore Health Services Institutional Animal Care and Use Committee (Singhealth IACUC 2012/SHS/757). Moreover, the procedures also comply to the recommendations prescribed in [32]. The animals were implanted with a titanium head post (Crist Instruments, MD, USA) prior to implantation of microelectrode arrays (MicroProbes, MD, USA - NHP A; Blackrock, UT, USA - NHP B) in the hand/arm region of primary motor cortex.

### A. Signal Acquisition and Processing

#### 1) Experiment 1 (NHP A)

4 microelectrode arrays containing 16 electrodes each were implanted in the hand/arm region of the left primary motor cortex in NHP A. A 100-channel neural recording system [33], [34] was used for acquiring neural data sampled at 13 *k*Hz from the implanted 64 electrodes. This raw data was then digitally band-pass filtered in MATLAB between 300 and 3000 Hz [19]. Typically, absolute threshold based method [35] is employed for the purpose of spike detection on commercial recording equipments. However, for this custom recording setup [33], [34], we found non-linear energy operator (NEO) based spike detection method [36] to be more effective over AT, and hence we used NEO to isolate spikes. The probable reason for this is the increased input-referred noise in custom recording setups compared to the commercial ones [37], [38]. First 30 seconds of filtered raw data *x*[*n*] on each day was used to set the spike detection threshold as,

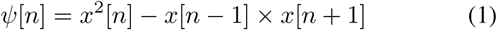

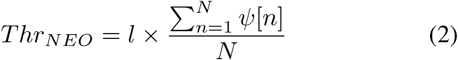

where *ψ*[*n*] is the NEO processed, pre-emphasized signal at time *n. Thr*_*NEO*_ is then computed as a multiple - *l* of average of *ψ*[*n*] over *N* samples of filtered raw data. The optimal value of *l* is found to be 13. Threshold was computed at the beginning of each day and was held fixed for the entire day’s analysis. NEO processed filtered raw data surpassing the threshold were classified as spikes and were used for subsequent analysis.

#### 2) Experiment 2 (NHP B)

1 microelectrode array containing 100 electrodes was implanted in the hand/arm region of the left primary motor cortex in NHP B. Ripple (UT, USA) grapevine neural interface processor was used for recording raw data sampled at 30 *k*Hz from two front-ends capable of recording 32 channels each. Ripple front-end processor provided bandpass filtered data between 250 and 7500 Hz to yield *x*[*n*] [39]. First 10 seconds of each day’s recording was used to compute threshold following absolute threshold method [35] as,

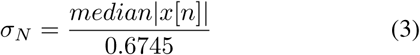

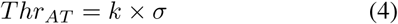

where *σ*_*N*_ is the standard deviation estimate over *N* samples of filtered raw data *x*[*n*], *k* is an integer typically between 2 and 5. For this dataset, we have employed *k* = 3. Spikes were computed as negative threshold crossings.

### B. Behavioural Tasks

The experiments involved an NHP trained to maneuver a joystick to reach a visual target. The joystick was spring-loaded to return to the centre position when released. Furthermore, it was restricted to move only in forward, right and left directions. The joystick position was tracked every 100 ms, and the instantaneous position of joystick was used to drive an effector in either of the three aforementioned directions or remain stationary. Appropriate joystick position thresholds were defined in each direction such that the effector moved once the position threshold was exceeded.

#### 1) Experiment 1 (NHP A)

This experiment has been described in detail in [19]. Briefly, the experiment consisted of four tasks - a) moving forward by 2 m, b) turning 90° right, c) turning 90° left and d) staying still for 5 seconds (*stop* task).

A trainer holding a piece of fruit reward served as a visual target for NHP A seated in a robotic wheelchair. A movement related trial was considered successful and rewarded with a piece of fruit, if the NHP reached the trainer within 15 seconds. For the stop trial, a task was considered successful and rewarded with a piece of fruit, if the monkey stayed still for a period of 5 seconds. The control of the robotic wheelchair was turned on at the beginning of each trial and was automatically turned off when it reached the target.

#### 2) Experiment 2 (NHP B)

In this experiment, a target is presented in a pseudo-random manner in one of the three different locations – Top, right and left relative to the centre of the screen in the form of a red square of side 2 cm on a black background. NHP B is seated in front of the screen in a primate chair. It is trained to drive the wheelchair avatar from the centre of the screen towards the target area. Once the wheelchair reaches the target area, the target’s colour changes from red to green and it serves as a cue for the NHP to stop moving. If the NHP manages to reach and stay in the target area under a total elapsed time of 13 seconds, the trial is considered successful and a juice reward is dispensed. The wheelchair avatar is driven at a frequency of 10 Hz in a manner similar to experiment 1, wherein instantaneous joystick signals are tracked to update the position of the wheelchair avatar.

## III. Proposed Decoding Algorithm

### A. Analysis Methodology

Incoming spike firing rates at time *t* = *i* denoted as *r*_*z*_(*i*) are computed at each of the *D* electrodes as the number of spikes occurring at a particular electrode *z* looking back in a time window *T*_*w*_ = 500 ms to yield input feature vector 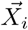 as,

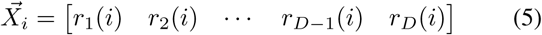

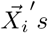 are computed every *T*_*s*_ = 100 ms for the entire duration of a trial. Correspondingly, joystick signals are tracked every *T*_*s*_ to yield the target value represented as a one-hot encoded vector - 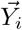 at time *t* = *i*. Target value can correspond one of the four discrete values of *forward, right, left* and *stop* in our reported experiments. Accumulating 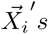 and 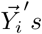 over all successful trials across all training sessions yields training matrix **X** and training target **Y**, where **X** *ϵ* ℝ^*D*×*n*^, **Y** *ϵ* ℝ^*C*×*n*^, *n* is the accumulated total number of instances across all training sessions and *C* is number of possible discrete target values. Only successful trials are used in this analysis. We used all testing data instances corresponding to successful trials in both NHPs. In NHP A, there was no problem of class imbalance, so we used all the training data instances corresponding to the successful trials. In NHP B, owing to the stop condition in each trial, there was a class imbalance problem. Hence, we employed an equal number of training data instances belonging to each class *forward, right, left* and *stop* sampled randomly on each day. Training a decoder involves learning a mapping between **X** and **Y**. Thus, a decoder outputs one of the four discrete outputs of *forward, right, left* and *stop* every *T*_*s*_ = 100 ms.

We have used a total of 35 sessions across 8 days recorded over a period of 66 days for NHP A. Typically, each day consists of 3 − 5 sessions and each session consists of 20 successful trials. Similarly, for NHP B, we have used a total of 30 sessions across 10 days recorded over a period of 77 days. For NHP B, we recorded 3 sessions every day with each session consisting of 60 successful trials. The number of input channels (*D*) correspond to 64 in both NHPs A and B.

In Fig. 4, we present the methodology of partitioning of data to be used as training and testing sets. We train fixed models on days 1 − 3 for both datasets and compare them against daily retrained models on the remaining days. The training days span a period of 12 and 17 days for NHP A and B respectively. In case of daily retraining, we have used 5 trials of each type in NHPs A and B. One must note that the test data for evaluating both types of models is the same.

**Fig. 4.**
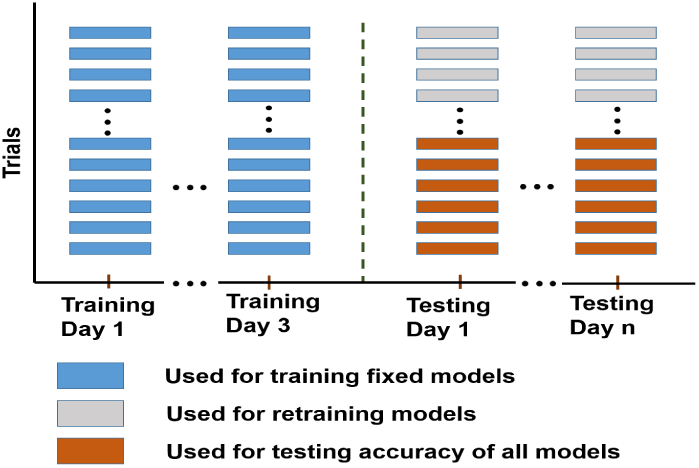
Partitioning of data for training and testing to benchmark fixed models against daily retrained models

### B. Sparse Ensemble Machine Learning

Ensemble methods employ a divide and conquer approach, wherein a set of classifiers attempt to independently learn a hypothesis each, which when combined effectively translates into a more robust classifier [40], [41]. In our case, the ensemble is made up of *M* base learners as shown in Fig. 5. The recording channels exhibit a certain degree of variability in firing rates across days. Micro-motion of electrodes as shown in Fig. 1 are a probable reason for this variability as neuronal signals appear/disappear at the electrodes. Thus, each of the classifiers in the ensemble are trained on a different set of input recording channels - 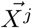, sampled with replacement from the entire set 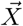, thereby reducing bias and contributing towards robustness [42]. The channels in each set are chosen randomly and the optimal number of channels (*U*) in each set are computed as,

**Fig. 5.**
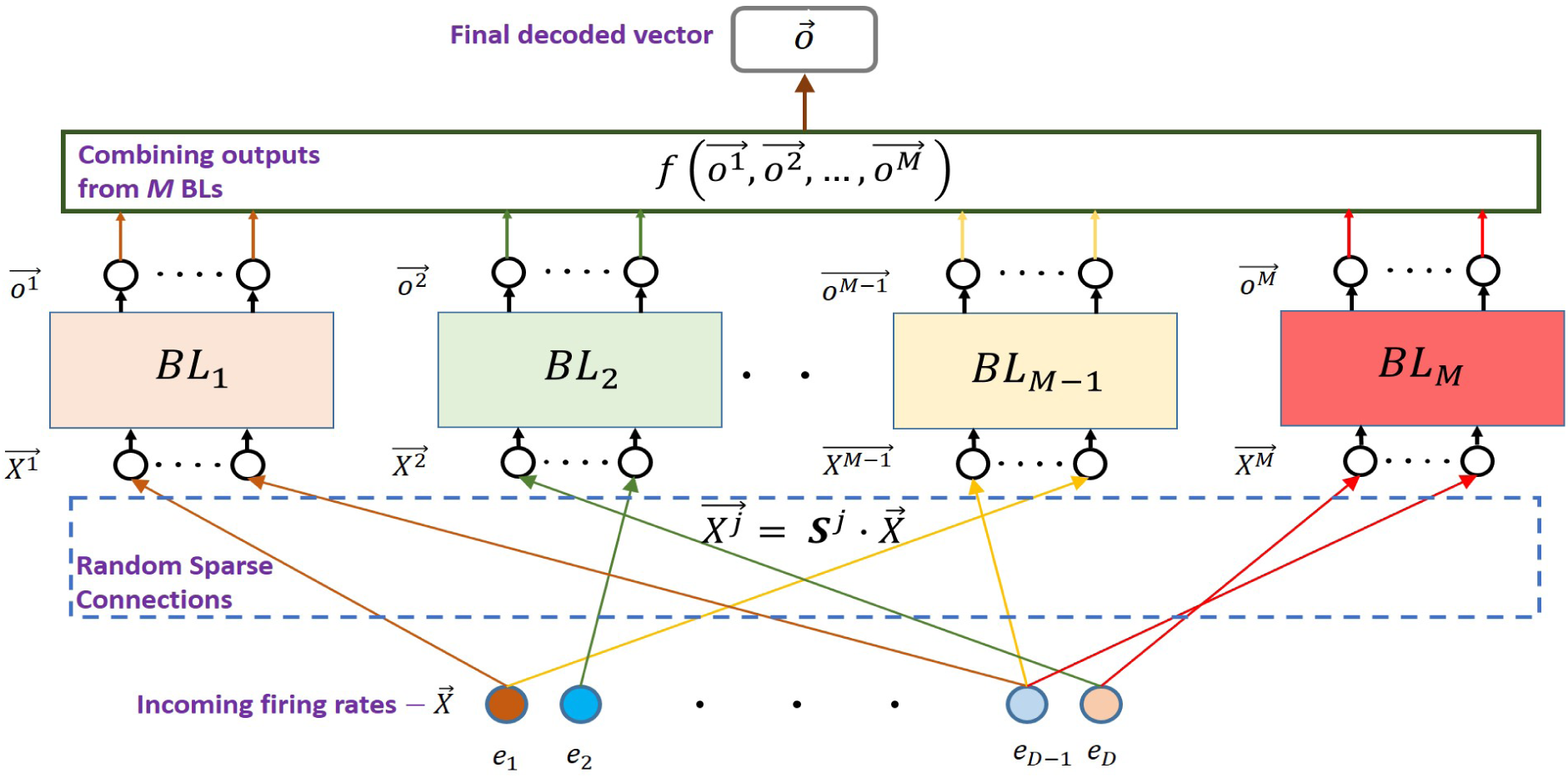
Schematic representation of sparse ensemble machine learning is presented here. Input to each of the *M* Base Learners (BLs) -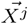 is obtained by sparsely sampling entire input space -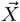 with replacement. Outputs 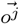 from all *M* constituent BLs are combined to obtain the final decoded vector 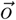

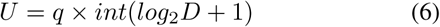

based on the suggestion imparted by Leo Breiman in his seminal Random Forest paper [43]. *D* as stated previously represents the total number of input channels and *q* is an integer hyper-parameter. The number of channels chosen in each set dictates the strength of each classifier and the correlation between them. A certain degree of heterogeneity is desired among the different sets [40], [41], [43].

Mathematically, 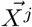 can be arrived at as follows,

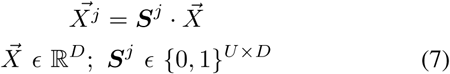

where ***S***^*j*^ is a sparse selection matrix. In this work, we restrict ***S***^*j*^ to have exactly one 1 in every row and column in a random location. This amounts to selecting a random subset of electrodes as input 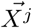 to *j*^*th*^ base learner. The number of such unique matrices is given by 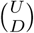.

Finally, the output 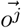 from each of the *M* base learners needs to be combined to yield a final output 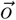.

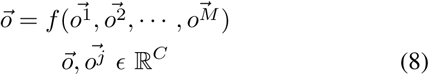

In this work, we have explored two intuitive ways to arrive at the final output as outlined below.

#### 1) Method 1

We simply sum the outputs from all *M* base learners to obtain 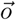 as shown below,

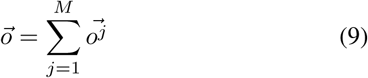

Final decoded value corresponds to the output class of 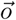 having the highest value among all *C* classes.

#### 2) Method 2

An alternative method is to perform voting among the decision taken by each of the *M* base learners,

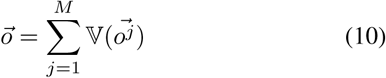

where 𝕍: ℝ^*C*^ *→* {0, 1}^*C*^ is a function whose output is 1 for the output class of 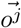 having the maximum value. Final decoded value in this case corresponds to the output class of 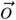 having maximum score among *C* classes.

## VI. Results

We present decoding comparison of two variants of fixed models - a) a single classifier based model trained on the entire feature set and b) a sparse ensemble classification model - in order to highlight the effectiveness of our proposed approach. These fixed models are trained on the initial 3 days as shown in Fig. 4. Linear discriminant analysis (LDA), support vector machine (SVM), extreme learning machine (ELM) and multilayer perceptron (MLP) have been used to compare the two variants of fixed models. Furthermore, we also contrast the decoding results of fixed models against a daily retrained model.

LDA and SVM have been quite popularly employed in both invasive and non-invasive forms of brain machine interfaces [44]–[46]. ELM has been shown to exhibit comparable results to the state of the art algorithms such as SVM, along with the benefit of being trained quickly on a small amount of training data [47], [48]. MLPs and its variants have been enjoying popular support with the dawn of the deep learning revolution and have recently shown impressive results in iBMIs [49]. The aforementioned reasons drove us towards choosing these algorithms as a means of providing a comprehensive benchmark for fixed classification models and their sparse ensemble versions. For a daily retrained model, simple linear models have been shown to perform reasonably well with small amount of training data [19], [24] and hence we have used LDA to obtain a daily retrained model.

### A. Choice of Hyperparameters

Sparse ensemble decoder training involves optimal tuning of - number of inputs to each base learner (*U*) and the number of base learners (*M*). In general, we expect the decoding accuracy to increase as we increase *M* at the cost of increased computation. However, the relation with *U* is not so obvious and we explore that first. We employ the well known k-fold cross-validation method for this purpose [50].

*U* is set as per Equation 6 and the decoding accuracy is observed by sweeping *q* as shown in Fig. 6 for NHP A and B’s data respectively. This is done first while sweeping *M* from 10 to 25 in steps of 5. However, it is found that regardless of the value of *M*, decoding accuracy peaks for *q* = 2 for NHP A and *q* = 5 for NHP B across all algorithms. The corresponding optimal value of *U* comes out to be 14 (*q* = 2) and 35 (*q* = 5) for NHP A and NHP B respectively.

**Fig. 6.**
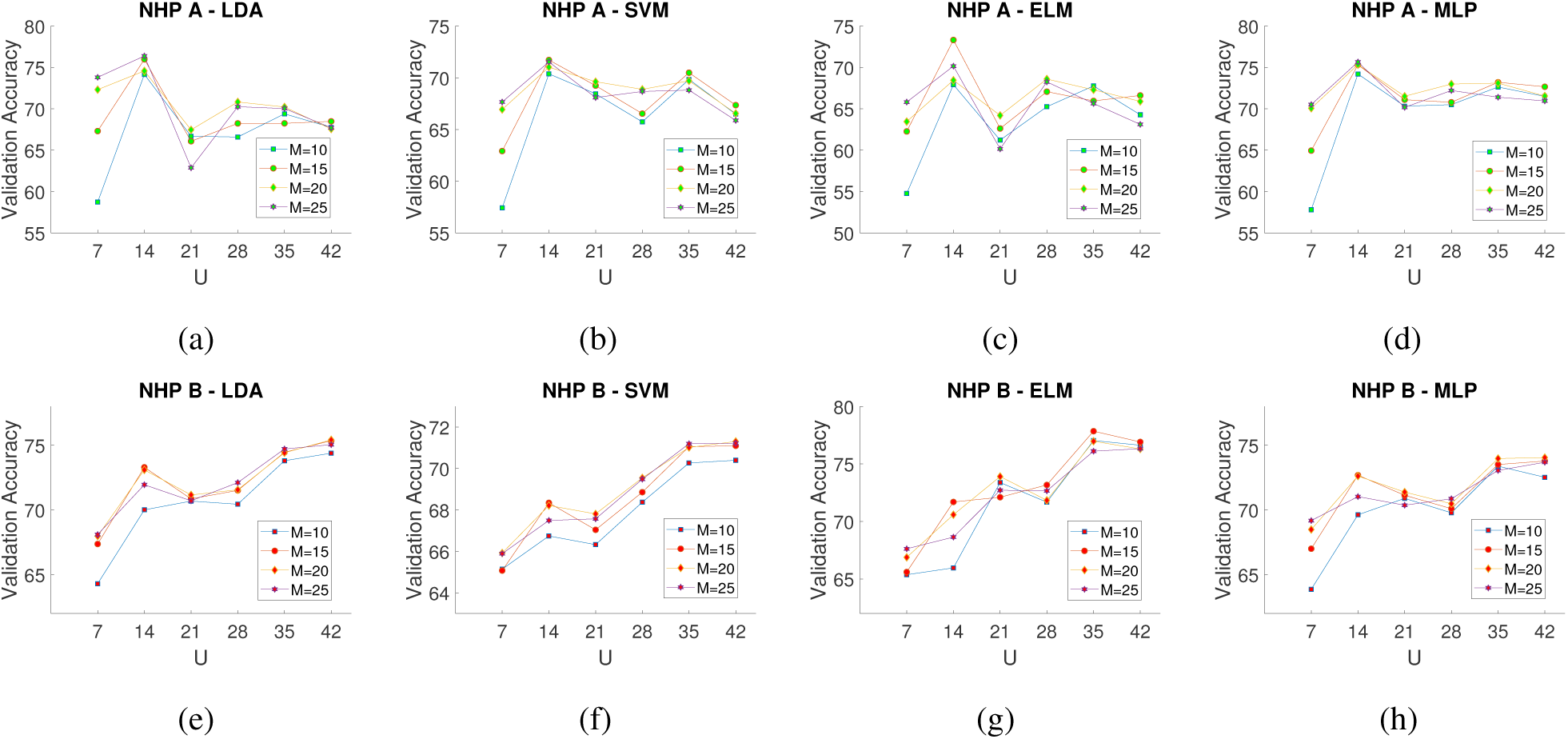
k-fold cross-validation to obtain optimal value of number of inputs (*U*) to each base learner as per Equation (6) for NHP A - (a),(b),(c),(d) and NHP B - (e),(f),(g),(h). The results are obtained for different number of base learners (*M*) swept from 10 to 25 across different classifiers. The optimal value of *U* comes out to be 14 (*q* = 2) and 35 (*q* = 5) across classifiers for NHP A and NHP B respectively.

Similarly, the optimal number of base learners, *M*, is next determined by sweeping its value (in a much finer grid than in Fig. 6) as shown in Fig. 7, while choosing optimal *q* found earlier. The optimal value is found to be around *M* = 13 across algorithms for both cases which is close to the value of *M* = 15 obtained in the coarse sweep of *M* in earlier experiments.

**Fig. 7.**
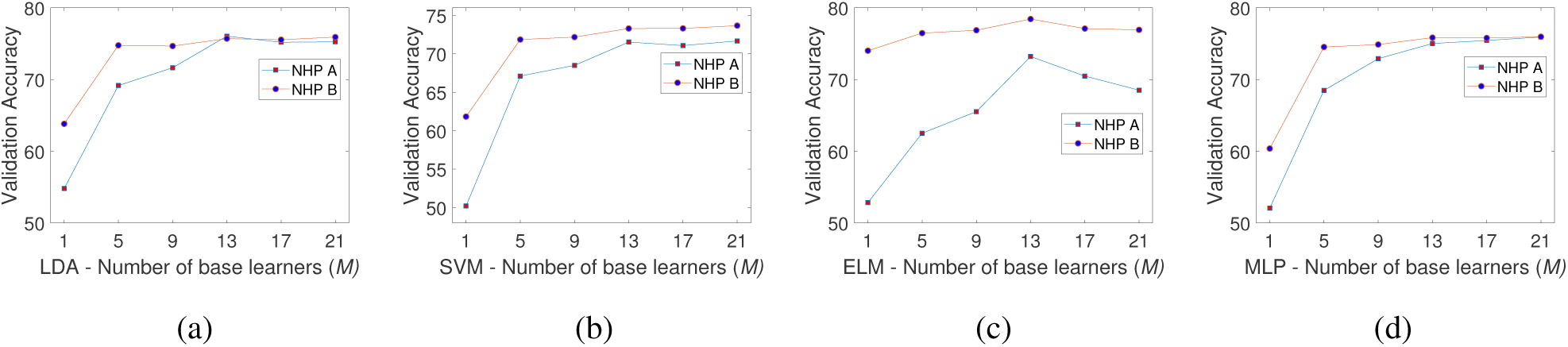
k-fold cross-validation to obtain optimal value of number of base learners (*M*) for NHP A and NHP B across different classifiers for the value of *U* obtained in Fig. 6. The optimal value of *M* comes out to be around 13 for both NHPs.

### B. Base Learners training

LDA does not require any hyperparameter tuning. ELM requires only tuning of number of hidden layer neurons and *k* − *fold* cross-validation was used to arrive at an optimal value. No activation function was employed at the hidden nodes in ELM. We used SVM with a Gaussian kernel, and Bayesian optimization [51] was employed to automatically tune *BoxConstraint* and *KernelScale* hyperparameters in a *k* − *fold* cross-validation regime. MATLAB2019a (The MathWorks, Inc., Natick, Massachusetts, United States) function *bayesopt* was used over 30 iterations to arrive at the optimal model. We have used a two hidden layer MLP with bayesian optimization employed for hyperparameter tuning [52]. *Hyperopt* [53] library was used to tune the following hyperparameters listed in Table. I over 200 evaluations.

**TABLE I.**
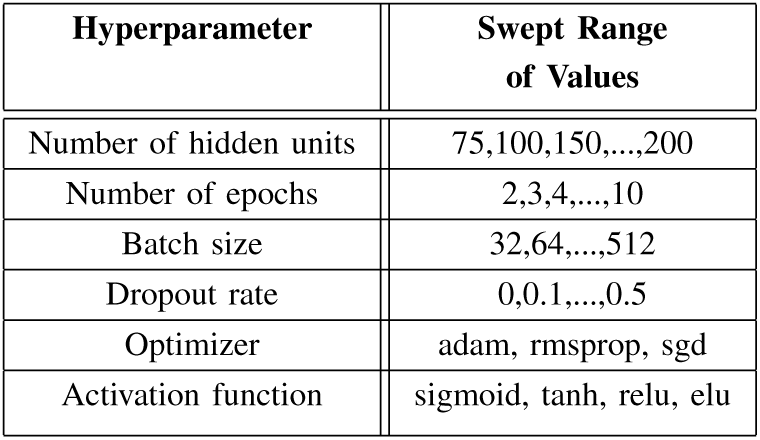
Hyperparameter Tuning for a Two Hidden Layer MLP Using Hyperopt Library.

Table. II gives an estimate of the training time involved in training a single classifier model for the different algorithms on NHP A’s dataset. *fitcdiscr* and *fitcecoc* functions defined in MATLAB2019a were used for training LDA and SVM models respectively. ELM was implemented in MATLAB2019a. MLP was implemented in *Python*3 on a jupyter notebook using Keras library [54]. Simulations were run on an Intel Core i5 powered laptop to report the above timings. One can observe that for our dataset, complex algorithms - SVM and MLP take ≈ 4 orders of magnitude of more time to train compared to single shot learning algorithms - LDA and ELM.

**TABLE II.** Training Time for Unoptimized Code of Single Classifier Models for NHP A.

| Algorithm | Training Time (in seconds) |
| --- | --- |
| LDA | 0.15 |
| SVM | 2438.4 |
| ELM | 0.08 |
| MLP | 2761.3 |

### C. Decoding Results

With the hyperparameters fixed based on initial days’ training data, we now evaluate the robustness of decoding over multiple days of neural recording in two versions of test datasets. The first version is the originally recorded test dataset referred to as the *unperturbed* dataset. In the second version, we have dropped the most informative electrode from the test day’s data by setting its firing rate to zero as a way of increasing the severity of nonstationarity [17]. This test dataset is referred to as the *perturbed* dataset. Most informative electrode was computed as the electrode having the highest mutual information score between firing rate **X** and target **Y** [55].

The *unperturbed* test results for our proposed sparse ensemble method against the corresponding single classifier models across different algorithms can be seen in Fig. 8. Maximum improvements of 20.82%, 12.92%, 18.66% and 9.53% in NHP A and 7.15%, 8.65%, 6.73% and 8.71% in NHP B are obtained by sparse ensemble methods over single classifier counterparts across LDA, SVM, ELM and MLP algorithms respectively. Furthermore, average improvements of 11.67±6.39, 7.51±3.37, 9.01±6.74, 4.73±4.16 in NHP A and 3.16 ± 2.66, 3.03 ± 2.79, 3.13 ± 1.73, 3.83 ± 3.86 in NHP B are obtained by sparse ensemble methods for LDA, SVM, ELM and MLP algorithms respectively.

**Fig. 8.**
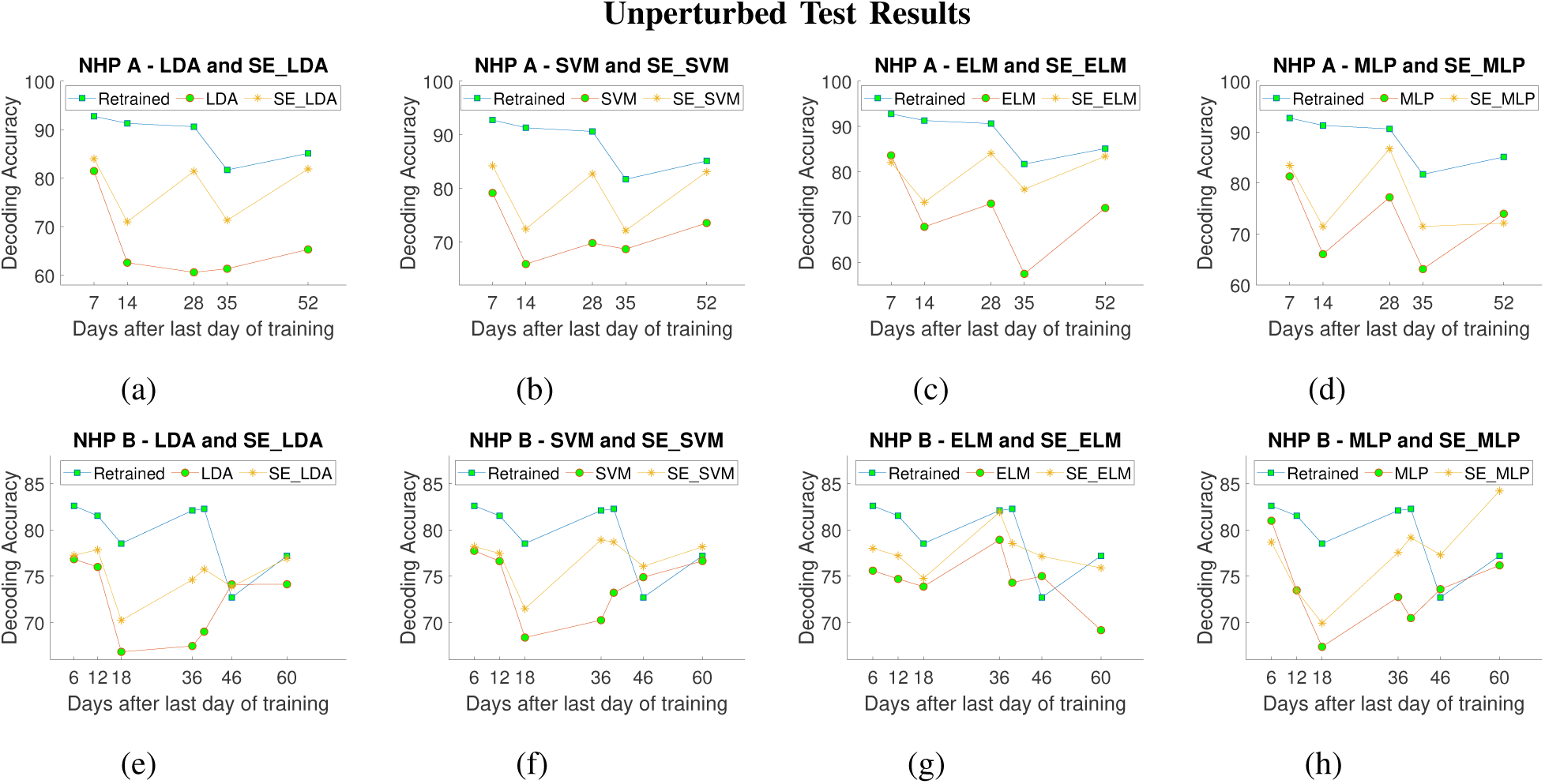
Decoder results across days for NHP A - (a),(b),(c),(d) and NHP B - (e),(f),(g),(h) have been presented. Classification models and their **<u>S</u>**parse **<u>E</u>**nsemble (SE) versions have been compared alongside a daily retrained model in the above plots. Outputs from constituent base learners are simply summed to arrive at the final output in SE models in the above reported case.

Decoding accuracies for *perturbed* test sets are reported in Fig. 9. Maximum improvements of 24.16%, 11.44%, 21.66% and 8.82% in NHP A and 13.76%, 18.96%, 7.33% and 28.42% in NHP B are obtained by sparse ensemble methods over single classifier counterparts across LDA, SVM, ELM and MLP algorithms respectively for perturbed test sets. Furthermore, average improvements of 12.33 ± 6.60, 7.11 ± 2.89, 9.21±8.64, 3.68±3.76 in NHP A and 8.27±4.28, 13.00±5.48, 3.73 ± 2.38, 14.08 ± 7.12 in NHP B are obtained by sparse ensemble methods for LDA, SVM, ELM and MLP algorithms respectively.

**Fig. 9.**
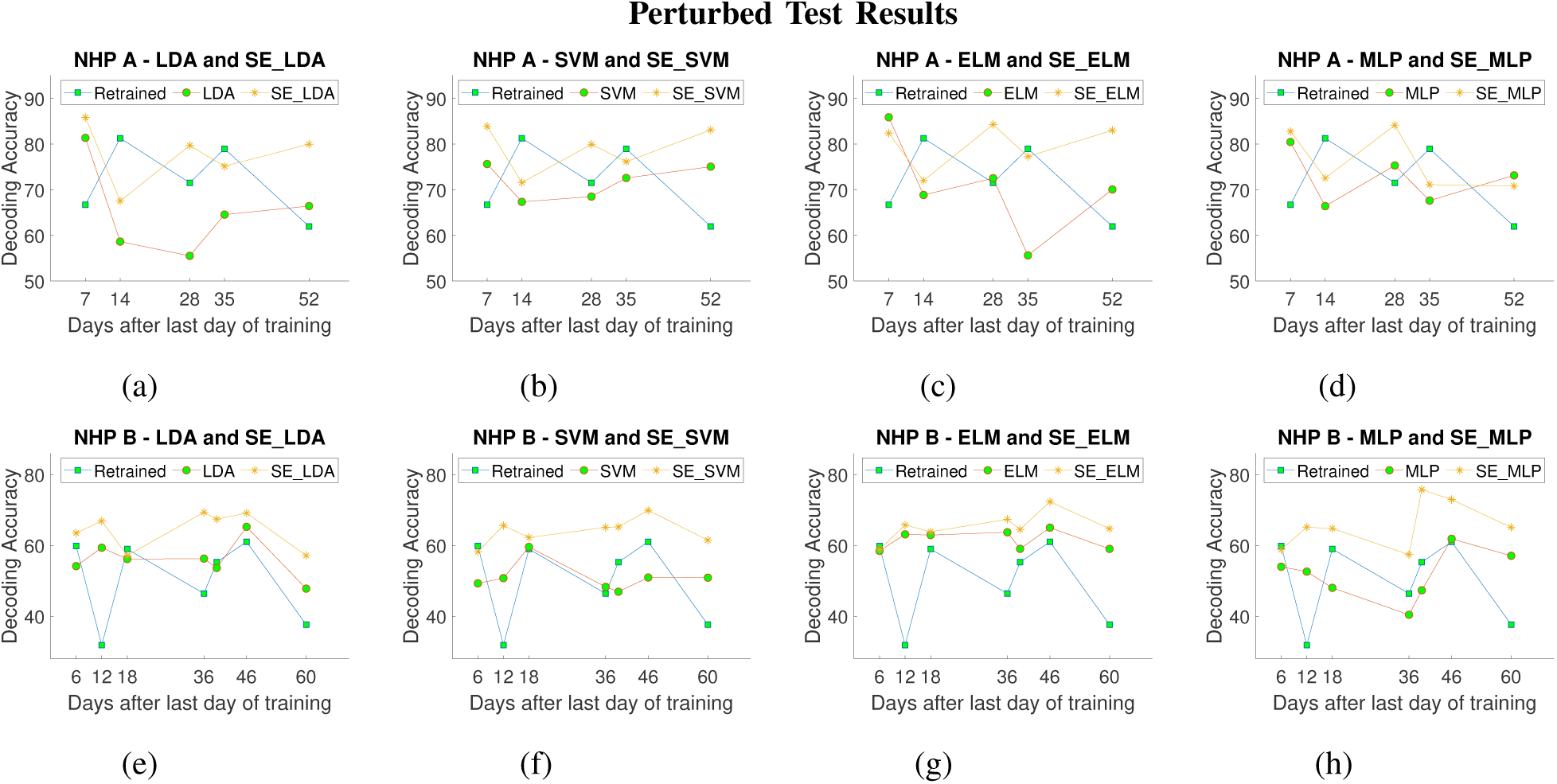
Decoder results across days for NHP A - (a),(b),(c),(d) and NHP B - (e),(f),(g),(h) while dropping the most informative electrode on the test day are presented. Fixed classification models and their **<u>S</u>**parse **<u>E</u>**nsemble (SE) versions have been compared alongside a daily retrained model in the above plots. Outputs from constituent base learners are simply summed to arrive at the final output in SE models in the above reported case.

Statistical comparison of all the single classifier test day results comprising of LDA, SVM, ELM and MLP algorithms was performed against their sparse ensemble versions for a two-tailed Wilcoxon signed-rank test. Significance values of 2.93 × 10^−4^, 1.43 × 10^−4^, 7.79 × 10^−4^ and 3.79 × 10^−6^ are obtained for NHP A-unperturbed, NHP B-unperturbed, NHP A-perturbed and NHP B-perturbed cases respectively. Thus, statistically significant results (*p* < 0.01) are obtained when using sparse ensemble methods over a single classifier.

A baseline daily retrained LDA based classifier outperforms fixed sparse ensemble and single classifier methods in line with similar results presented in [24], across both NHPs in the unperturbed case in Fig. 8. In Fig. 9, we present results of a retrained LDA model on held out test trials (Fig. 4) while dropping the most informative electrode in held out test trials only. As expected, a drastic drop is observed in the daily retrained model. In fact, sparse ensemble versions of LDA, SVM, ELM and MLP outperform a daily retrained model by an average 5.54%, 6.83%, 7.70%, 4.18% in NHP A and 14.21%, 13.81%, 15.20%, 15.55% in NHP B respectively.

## V. Discussion

### A. Combining Strategy: Sum vs Voting

As shown in Section III-B2 voting as a method of combining outputs is also quite popular. However, we have presented detailed results using the summing approach since it consistently outperformed the voting approach as seen in Fig. 10 across classifiers - LDA, SVM, ELM and MLP.

**Fig. 10.**
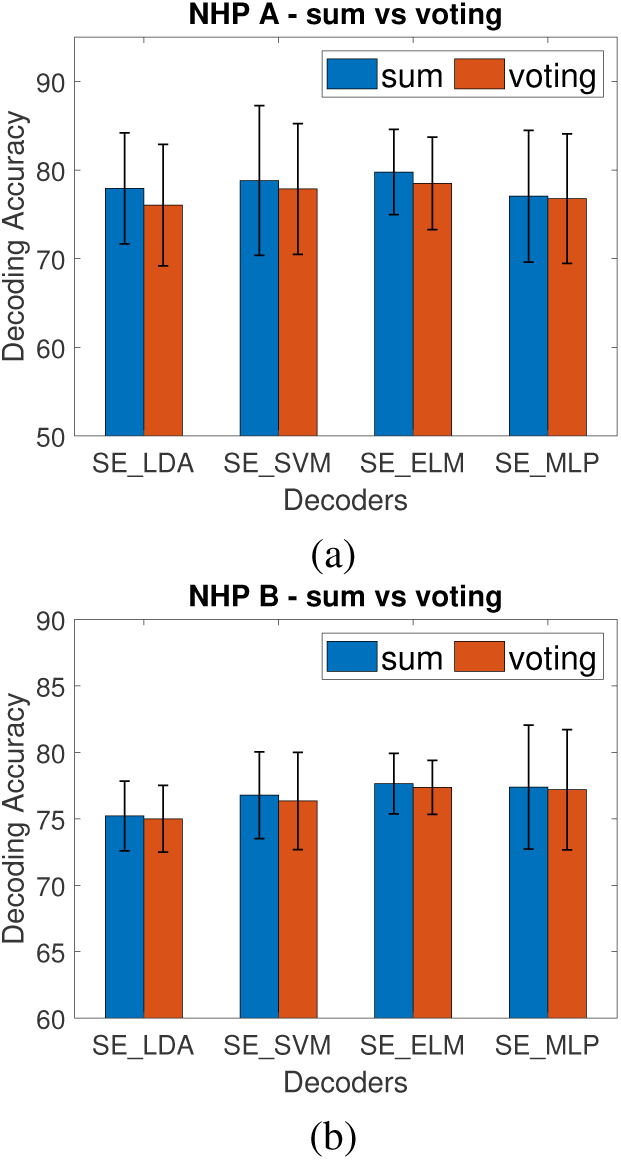
Average decoded results across test days for NHPs A and B in (a) and (b) respectively comparing summation and voting methods to arrive at the final ensemble output across different methods.

### B. Ensemble vs Sparse Ensemble

To validate if *sparsely* sampling input electrodes to the individual base learners is indeed responsible for robustness against simply forming ensembles with all input features, we present comparison of *SE*_*ELM*_*sum* against *Ensemble*_*ELM*_*sum* for NHP A in an unperturbed setting in Fig. 11. We have used summation as method of combining outputs in both methods with hyperparameter tuning carried out as per the methodology detailed in *Section* IV-A. ELM was chosen for comparison as it is seen to yield a performance comparable to the other classifiers in both NHPs. NHP A exhibits relatively more nonstationarity compared to NHP B as seen in Fig. 3 and hence we have used it in the below example. From Fig. 11, we can see that *SE*_*ELM*_*sum* outperforms *Ensemble*_*ELM*_*sum* by an average 11.78% highlighting the important role played by the sparse connection of input electrodes to each base learner.

**Fig. 11.**
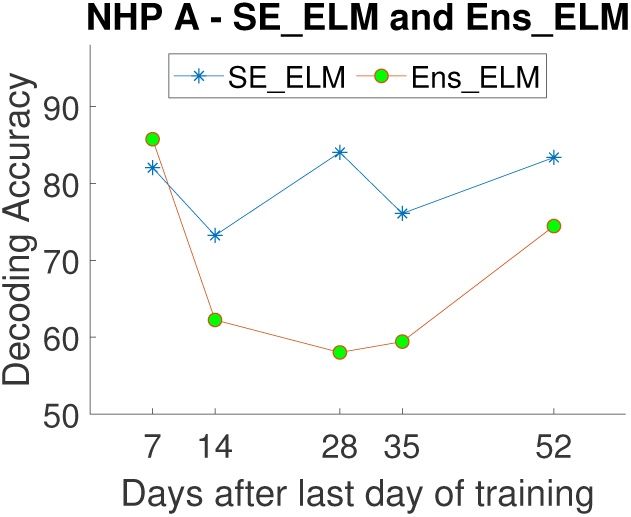
Comparing Sparse Ensemble ELM (*SE_ELM*) against Ensemble ELM (*Ens_ELM*) with summation as a method of combining outputs in unperturbed NHP A dataset. This figure highlights that introducing sparsity in the inputs to base learners lends robustness to ensemble based learning.

### C. Comparison

[17] has reported steady improvements in offline decoding results over a daily retrained decoder in two NHPs spanning roughly two months. However, we believe approaches in general advocating for employment of complex deep neural networks [17], [28] to counter nonstationarity suffer from mainly three problems. Firstly, they seem to require large amounts of heterogeneous data spanning over a period of upwards of a year or more to perform at their best. Secondly, the number of hyperparameters and their subsequent tuning for optimal performance is a serious impediment to be overcome in these complicated neural networks [53], [56]. Thirdly, they risk running into latency issues owing to its computational complexity in terms of the number of operations that need to performed to arrive at the decoded output [28], [57]. Furthermore, other problems such as longer training times, interpretability and ease of use prevail [58].

It has been shown with theoretical support that ensemble based systems tailored to alleviate the source of nonstationarity can lend to more stable results compared to single classifier based systems [59], [60]. Based on this reasoning, we chose to build an ensemble of classifiers, with each classifier being trained on a randomly chosen subset of features with replacement [42]. The intuition behind this approach is that multiple hypotheses corresponding to different subsets of electrodes would be learned by individual classifiers in the ensemble. Thus, with the introduction of appropriate diversity and method of aggregating the outputs, the ensemble would be robust against incoming nonstationarity. On the contrary, a single classifier based system trained on a given training data distribution tends to fail with a differing test data distribution [61].

### D. Custom neural recording tools

Design of custom miniaturized neural recording setups such as [34] involve trade-offs between noise, power and area [37]. The quest for lower power and area consumption in miniaturized chips often drives up the value of input referred noise. For example, the custom recording setup [34] reports an input referred noise of 4.4 *µV RMS*, whereas commercial front ends such as Ripple [39] report an input referred noise levels of less than 2.1 *µV RMS*. Thus, we hypothesize that this increased recorded noise in a custom setup for NHP A is the reason for (a) increased sensitivity in selecting an optimal *M* (Fig. 7) and (b) need for pre-emphasis through NEO for effective spike detection. Despite this noise impediment in custom recording apparatus, sparse ensemble methods have the potential to yield consistent improvements as seen in the *Results* section.

## VI. Conclusion and Limitations

In this study, we have presented a novel way to increase the robustness of decoding over a longer term in iBMIs. This is the first ever study reporting consistent improvement against state of the art techniques over a period spanning roughly two months in two NHPs. Our novel and simple approach tackles the root of the nonstationarity problem, i.e. randomly changing firing rates and accounts for these changes by dividing a subset of the inputs randomly among each of the learners in an ensemble. Significant improvements can be seen in both actually recorded and manually perturbed data where the perturbation removes the most informative electrode(Fig. 8 and Fig. 9 respectively).

The efficacy of the sparse ensemble based approach remains to be validated in a closed-loop setting. Another possible limitation one might refer to is the simplistic nature of the reported experiment with relatively lower speeds of 10 Hz and a 4-options discrete control. However, one must note that despite its simplistic nature, the phenomenon of nonstationarity is at work and significant performance degradation is observed in fixed decoder models in both NHPs, particularly more so in NHP A. Do simple classifiers such as LDA, ELM as constituent base learners suffice to yield a reasonable robust response or as more data is eventually available with iBMI use, does it help to switch to more complex base learners is an interesting strand of research we aim to pursue in the near future.

## Acknowledgment

The authors would like to thank Abdur Rauf and Clement Lim for helping in training the NHPs and data collection. This work was supported through grant RG87/16 by MOE, Singapore.

